# Unveiling effects of growth conditions on crown architecture and growth potential of Scots pine trees

**DOI:** 10.1101/2021.12.13.472374

**Authors:** Ninni Saarinen, Ville Kankare, Saija Huuskonen, Jari Hynynen, Simone Bianchi, Tuomas Yrttimaa, Ville Luoma, Samuli Junttila, Markus Holopainen, Juha Hyyppä, Mikko Vastaranta

## Abstract

Trees adapt to their growing conditions by regulating the sizes of their parts and their relationships. For example, removal or death of adjacent trees increases the growing space and the amount of light received by the remaining trees enabling their crowns to expand. Knowledge about the effects of silvicultural practices on crown size and shape as well as about the quality of branches affecting the shape of a crown is, however, still limited. Thus, the aim was to study the crown structure of individual Scots pine trees in forest stands with varying stem densities due to past forest management practices. Furthermore, we wanted to understand how crown and stem attributes as well as tree growth affects stem area at the height of maximum crown diameter (SAHMC), which could be used as a proxy for tree growth potential. We used terrestrial laser scanning (TLS) to generate attributes characterizing crown size and shape. The results showed that increasing stem density decreased Scots pine crown size. TLS provided more detailed attributes for crown characterization compared to traditional field measurements. Furthermore, decreasing stem density increased SAHMC and strong relationships (Spearman correlations >0.5) were found between SAHMC and crown and stem size as well as stem growth. Thus, this study provided quantitative and more comprehensive characterization of Scots pine crowns and their growth potential.

## Introduction

Trees direct available resources to reproduction and growth and can regulate their size and the relationship between their parts. That way trees adapt to changes in their growing conditions. The size of a tree correlates with the space a tree occupies and it defines tree growth which is linked to carbon sequestration (Pretzsch et al. 2015). Removal or death of trees enhances the light regime and photosynthesis for the remaining trees, which increases the crown size. This is particularly evident near the lowest limit of live crown where changes in the amount of light increases considerably more compared to the top of a tree (Oker-Blom & Kellomäki 1982).

Trees of different species require differing amount of growing space; birch (*Betula* sp.) requires more space than Scots pine (*Pinus sylvetris* L.) which in turn is more demanding than Norway spruce (*Picea abies* (H. Karst) L.) (Aaltonen 1925, Pretzsch et al. 2015). Additionally, crown architecture (e.g., crown width, live-crown length) varies between mixed stands compared to monocultures (Bauhus et al. 2004, Bayer et al. 2013, Dieler & Pretzsch 2013, Pretzsch 2014). There is a relationship between tree size and growing conditions that can be assessed through the light regime. In dense forests lower branches die due to the limited amount of light (Heikinheimo 1953, Flower-Ellis et al. 1976, Kellomäki 1980) specifically for light-demanding species such as Scots pines and birches (Kellomäki & Tuimala 1981), and this decreases live-crown ratio (i.e., proportion of live crown from tree height).

Forest management is mainly aimed at increasing size and quality of the trees left to grow by regulating stand density and thus improving their growing conditions. First commercial thinning is especially important for Scots pines and later thinnings, even if intensive, do not offer recovery from reduced live crown ratio as it has been shown to reduce up to 37% of tree height (Mäkinen & Isomäki 2004c). The crown of young trees recover better compared to old trees because height growth of young trees increases the length of live crown (Hynynen 1995). In mature and old trees, height growth is slower, and recovery of a crown is limited to increasing the width and the number of leaves/needles.

There is a long history of research where the relationship between crown and stem dimensions has been investigated (Krajicek et al. 1961, Larson 1963, Gingrich 1967, Curtin 1970, Seymore & Smith 1987). Process-based models simulate tree growth as a function of leaf biomass, in other words of their photosynthetic elements (e.g. Valentine & Makela, 2005). Shinozaki et al. (1964) proposed a conceptual framework for the relationship between the amount of stem tissue and corresponding supported leaves known as the pipe model theory (PMT). It has been shown that the total cross-sectional area of living branches correlated strongly with foliage mass (Vanninen et al., 1996; Ilomäki et al., 2003; Kantola & Mäkelä, 2005). Longuetaud et al. (2006) reported that statistically significant indicators for tree vitality were the total cross-sectional area of branches, height-diameter at breast height (DBH) ratio (i.e., height/DBH), and the relative and absolute height of the crown base. More specifically, Lehtonen et al. (2020) and Hu et al. (2020) found leaf biomass of Scots pine to be proportional to the stem cross sectional area at the crown base. However, in both cases the relationship was influenced by other factors, such as age, site type, and temperature. There are indeed criticisms on the validity of the PMT, for which we direct the reader to the extensive review from Lehnebach et al. (2018). In any case, if traditional empirical models are using DBH as a proxy for growth potential, the question still remains if diameter at the crown base (dcb) could be a more accurate predictor.

Crown attributes from standing trees have been limited to crown base height, crown length, live-crown ratio, projection area, and crown width of which the last one has been more challenging to measure from several directions. Laser scanning (or Light detecting and ranging LiDAR) has provided new opportunities for characterizing trees in more detail in three-dimensional space. Especially terrestrial laser scanning (TLS) has increasingly been used in producing a variety of tree attributes (Seidel et al. 2011, Metz et al. 2013, Seidel et al. 2015, Hess et al. 2018, Chianucci et al. 2020, Owen et al. 2020, Saarinen et al. 2017, Georgi et al. 2021, Rais et al. 2021, Zhu et al. 2021). One of the challenging stem-related attributes to be measured from standing trees has been taper curve (i.e., diameters at various heights of a stem) and TLS data has been shown to overcome that challenge (Liang et al. 2014, Yrttimaa et al. 2019, 2020). Additionally, versatile crown attributes such as volume (Fernández-Sarrá et al. 2013), surface area (Metz et al. 2013), asymmetry (Seidel et al. 2011), and height of the maximum crown projection area (Seidel et al. 2011) have been generated. Binkley et al. (2013) and Forrester (2014) have stated that crown projection area and crown volume, which can be obtained with TLS data, can be used as proxies for leaf area and leaf biomass. Furthermore, crown surface area has been used as a proxy for the photosynthetically active surface of the tree (Seidel et al 2019a). TLS has also been used for studying competition between species (Martin-Ducup et al. 2016, Barbeito et al. 2017, Juchheim et al. 2019, Pretzsch et al. 2019, Hildebrandt et al. 2021), the effects of management intensity on tree structure (Juchheim et al. 2017, Georgi et al. 2018, Bogdanovich et al. 2021), as well as structural complexity of individual trees (Seidel 2018, Seidel et al. 2019b, Saarinen et al. 2021). Thus, TLS provides a vast range of opportunities for understanding tree growth.

Knowledge about the effects of silvicultural practices on crown attributes such as volume and length as well as crown diameter and its variation that affect the shape of a crown is still limited. Thus, the aim is to investigate how crown structure of individual Scots pine trees varies when growing in differing conditions due to the intensity and type of past thinning treatments. It is hypothesized that crown size decreases with increasing stem density (H1) and increases when suppressed and co-dominant trees were removed (H2). Related to the PMT, the objective is to understand the relationship between stem area at the height of the maximum crown diameter (SAHMC) and crown and stem dimensions as well as growth of the tree. This relates to the question of the usefulness of diameter at the crown base (dcb) as a proxy for growth potential as it is of renewed importance since new technology such as TLS can now estimate this parameter more easily.

## Materials

The study area is located in southern boreal forest zone in Finland and consists of three study sites (Figure 1) with relatively flat terrain (elevation above sea level ~137 m±17 m) in mesic heath forest (i.e., Myrtillus forest site type according to Cajander (1909)) dominated by Scots pine.

**Figure 1.**
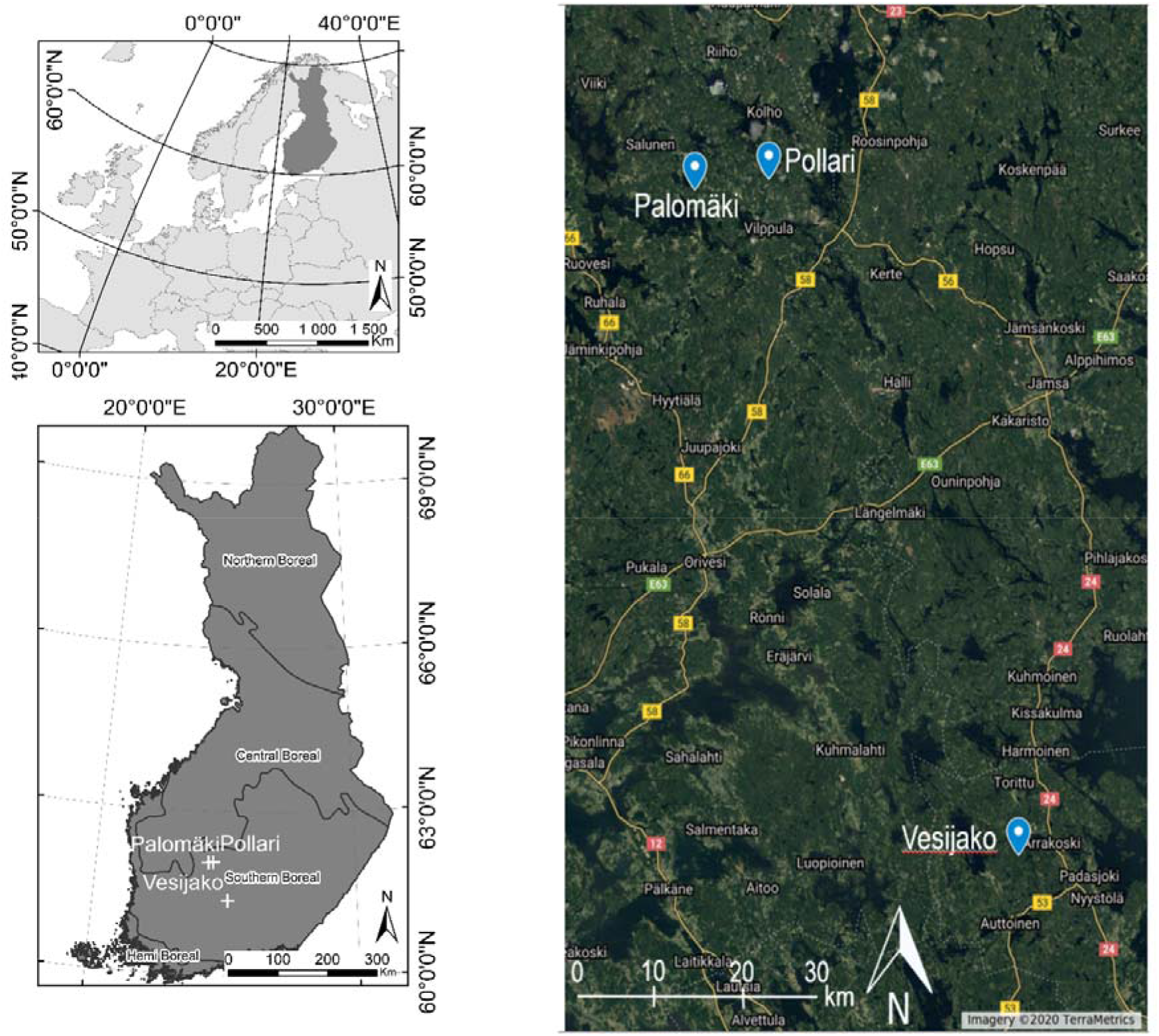
Location of tree study sites (i.e. Palomäki, Pollari, and Vesijako) and vegetation zones in Finland (bottom left) and study sites on top of satellite imagery © 2020 TerraMetrics.

The proportion of Norway spruce and deciduous trees (i.e., *Betula* sp and *Alnus* sp) from the total stem volume of all trees within the 27 sample plots was 3.06% and 0.03%, respectively. The study sites were established in 2005 and 2006 when nine rectangular sample plots (sized 1000-1200 m^2^) were placed on each study site. At the same time, first in situ measurements were carried out and the plots were also thinned according to the experimental study design that included two level of thinning intensity and three thinning types (Table 1). One plot at each study site was left as a control plot where no thinning has been carried out since the establishment of the sites. Thinning intensity was defined as the remaining basal area whereas thinning type determined which trees (based on a crown class) were removed. The remaining relative stand basal area after moderate thinning was ~68% of the stocking before thinning and intensive thinning reduced the stocking levels down to 34%. Suppressed and co-dominant trees were removed in thinning from below whereas dominant trees were mainly removed in thinning from above. Dominant trees were removed and small, suppressed trees were left to grow in systematic thinning from above without considering regular spatial distribution of the remaining trees, which was considered in thinnings from below and above. Additionally, unsound and damaged trees (e.g., crooked, forked) were removed in thinnings from below and above.

**Table 1.** Thinning treatments applied at the sample plots when the study sites were established.

| Thinning treatment | Acronym | Explanation | Number of plots | Number of stems / ha |
| --- | --- | --- | --- | --- |
| **Moderate thinning from below** | Moderate below | Moderate thinning refers to prevailing thinning guidelines applied in Finland (Rantala 2011) | 3 | 716 |
| **Moderate thinning from above** | Moderate above |  | 4 | 913 |
| **Moderate systematic thinning** | Moderate systematic |  | 5 | 938 |
| **Intensive thinning from below** | Intensive below | Intensive thinning corresponds 50% lower remaining basal area (m<sup>2</sup>/ha) than in the plots with moderate thinning intensity | 3 | 287 |
| **Intensive thinning from above** | Intensive above |  | 4 | 446 |
| **Intensive systematic thinning** | Intensive systematic |  | 5 | 466 |
| **Control** | No treatment | No thinning treatment since the establishment | 3 | 1245 |

Tree species, DBH from two perpendicular directions, crown layer, and health status were recorded for each tree within a plot during all in situ measurements (i.e., at the establishment, 10 years after the establishment, and between October 2018 and April 2019 for this study). Each sample plot also includes ~22 sample trees from which also tree height, live crown base height, and height of the lowest dead branch were measured. Plot-level attributes before and after thinning treatments (i.e. at the establishment) as well as based on the in-situ measurements in 2018-2019 are presented in Table 2, and the development of tree-level attributes for each thinning treatment can be found in Table 3.

**Table 2.** Mean and standard deviation (with ±) of stand characteristics by treatments at the establishment (2005-2006) and after the growth period (2018-2019). G = basal area, N = stem number per hectare, V = volume, D_w_ = mean diameter weighted by basal area, and H_w_ = mean height weighted by basal area.

| 2005-2006 |  |  |  |  |  |  |  |
| --- | --- | --- | --- | --- | --- | --- | --- |
|  | No treatment | Moderate below | Moderate above | Moderate systematic | Intensive below | Intensive above | Intensive systematic |
| G (m<sup>2</sup>/ha) | 27.6±6.7 | 18.3±2.1 | 18.5±1.1 | 18.2±1.1 | 8.9±0.8 | 9.1±0.8 | 8.7±0.7 |
| N/ha | 1336±97 | 719±130 | 955±258 | 988±129 | 292±55 | 479±113 | 522±183 |
| V (m<sup>3</sup>/ha) | 224.0±92.8 | 148.8±30.2 | 144.0±15.3 | 141.3±23.6 | 72.9±12.4 | 69.1±11.3 | 67.3±14.7 |
| D<sub>w</sub> (cm) | 17.8±3.4 | 18.7±2.4 | 16.9±1.9 | 16.5±1.6 | 20.4±2.7 | 16.5±2.5 | 15.7±3.0 |
| H<sub>w</sub> (m) | 16.1±3.3 | 16.5±1.9 | 15.7±1.2 | 15.6±2.1 | 16.9±1.8 | 15.3±1.9 | 15.5±2.7 |
| 2018-2019 |  |  |  |  |  |  |  |
|  | No treatment | Moderate below | Moderate above | Moderate systematic | Intensive below | Intensive above | Intensive systematic |
| G (m<sup>2</sup>/ha) | 37.1±4.6 | 28.4±2.5 | 28.3±28.3 | 27.6±1.8 | 15.9±0.7 | 16.1±1.2 | 15.9±1.6 |
| **N/ha** | 1249±159 | 705±113 | 915±214 | 938±111 | 287±65 | 446±82 | 466±172 |
| **V (m<sup>3</sup>/ha)** | 380.3±93.9 | 291.8±44.7 | 282.3±6.1 | 267.9±16.1 | 160.8±9.1 | 150.5±12.6 | 150.4±9.9 |
| **D<sub>w</sub> (cm)** | 21.2±3.0 | 23.5±2.2 | 21.2±1.9 | 20.7±1.2 | 27.5±3.1 | 22.3±2.1 | 22.2±3.0 |
| **H<sub>w</sub> (m)** | 21.3±3.1 | 21.7±2.0 | 21.0±1.1 | 20.3±1.4 | 21.6±1.6 | 19.5±1.2 | 20.0±2.2 |

**Table 3.** Mean tree-level attributes with their standard deviation (with ±) for each treatment at the year of the establishment (2005-2006) and after the growth period (2018-2019). DBH = diameter at breast height.

| **2005-2006** |  |  |  |  |  |  |  |
| --- | --- | --- | --- | --- | --- | --- | --- |
|  | **No treatment** | **Moderate below** | **Moderate above** | **Moderate systematic** | **Intensive below** | **Intensive above** | **Intensive systematic** |
| **DBH (cm)** | 15.4±4.6 | 17.6±3.3 | 15.3±3.3 | 14.8±3.5 | 19.3±3.4 | 15.1±3.1 | 14.8±4.1 |
| **Height (m)** | 14.7±2.6 | 15.9±1.9 | 15.3±1.2 | 14.6±1.9 | 16.5±1.8 | 14.8±1.8 | 14.7±2.6 |
| **Volume (dm<sup>3</sup>)** | 160.5±119.7 | 202.7±89.3 | 149.6±76.2 | 138.1±77.8 | 249.2±107.0 | 141.8±73.4 | 145.6±97.1 |
| **2018-2019** |  |  |  |  |  |  |  |
|  | **No treatment** | **Moderate below** | **Moderate above** | **Moderate systematic** | **Intensive below** | **Intensive above** | **Intensive systematic** |
| **DBH (cm)** | 18.7±5.0 | 22.2±3.7 | 19.3±4.3 | 18.8±4.2 | 26.4±3.9 | 21.1±3.5 | 20.8±4.3 |
| **Height (m)** | 20.2±3.0 | 21.2±2.1 | 20.4±1.6 | 19.4±2.2 | 21.2±1.7 | 19.1±1.5 | 19.6±2.8 |
| **Volume (dm<sup>3</sup>)** | 299.4±190.8 | 408.3±106.3 | 306.4±145.6 | 282.5±137.2 | 563.8±202.5 | 335.2±125.4 | 347.0±173.3 |

TLS data acquisition was carried out with a Trimble TX5 3D phase-shift laser scanner (Trimble Navigation Limited, USA) operating at a 1550 nm wavelength and measuring 976,000 points per second. This resulted in a hemispherical (300° vertical x 360° horizontal) point cloud with a point distance approximately 6.3 mm at 10-m distance. Eight scans were acquired from each sample plot between September and October 2018. Two scans were placed on two sides of the plot center and six auxiliary scans were placed closer to the plot borders (see Figure 1 in Saarinen et al. 2020). Artificial targets (i.e., white spheres with a diameter of 198 mm) were placed around each sample plot to be used as reference objects for registering the eight scans into a single, aligned coordinate system with a FARO Scene software (version 2018). The registration resulted in a mean distance error of 2.9 ± 1.2 mm, mean horizontal error was 1.3 ± 0.4 mm, and mean vertical error 2.3 ± 1.2 mm. LAStools software (Isenburg 2019) was used to normalize the point heights to heights above ground by applying a point cloud normalization workflow presented by Ritter et al. (2017).

## Methods

First, plot-level TLS point clouds were segmented to identify points from individual trees. Local maxima from canopy height models (CHMs) with a 20-cm resolution were identified using the Variable Window Filter approach (Popescu & Wynne 2004) and the Marker-Controlled Watershed Segmentation (Meyer & Beucher 1990) was applied to delineate crown segments. A point-in-polygon approach was applied for identifying all points belonging to each crown segment. To identify points that originated from stem and crown within each crown segment, a point cloud classification procedure by Yrttimaa et al (2020) was used. The classification of stem and non-stem points assumed that stem points have more planar, vertical, and cylindrical characteristics compared to non-stem points representing branches and foliage (Liang et al. 2012, Yrttimaa et al. 2020). The method by Yrttimaa et al. (2019, 2020) is an iterative procedure beginning from the base of a tree and proceeding towards treetop. More detailed description of the point cloud classification workflow can be found in Yrttimaa et al. (2019, 2020). The result of this step was 3D point clouds for each individual Scots pine tree (n = 2174) within the 27 sample plots.

We generated several attributes characterizing crown size and shape (Table 4). Points from TLS that were classified originating from branches and foliage (i.e., crown points) in the previous step were utilized. A 2D convex hull was fitted to envelope the crown points of each tree of which crown projection area was derived. Crown diameter, on the other hand, was defined as the distance between the two most outer points in xy-space of the 2D convex hull. To obtain crown volume and surface area, a 3D convex hull was fitted to the crown points. We also wanted to investigate crown shape and thus divided the crown points into height percentiles (i.e., slices) of 10% starting from the lowest points. Then, 2D convex hull was fitted for each slice and its area and diameter were similarly obtained to the maximum crown diameter. Furthermore, mean, standard deviation, and range (i.e., crown taper) of these slice diameters were saved.

**Table 4.**
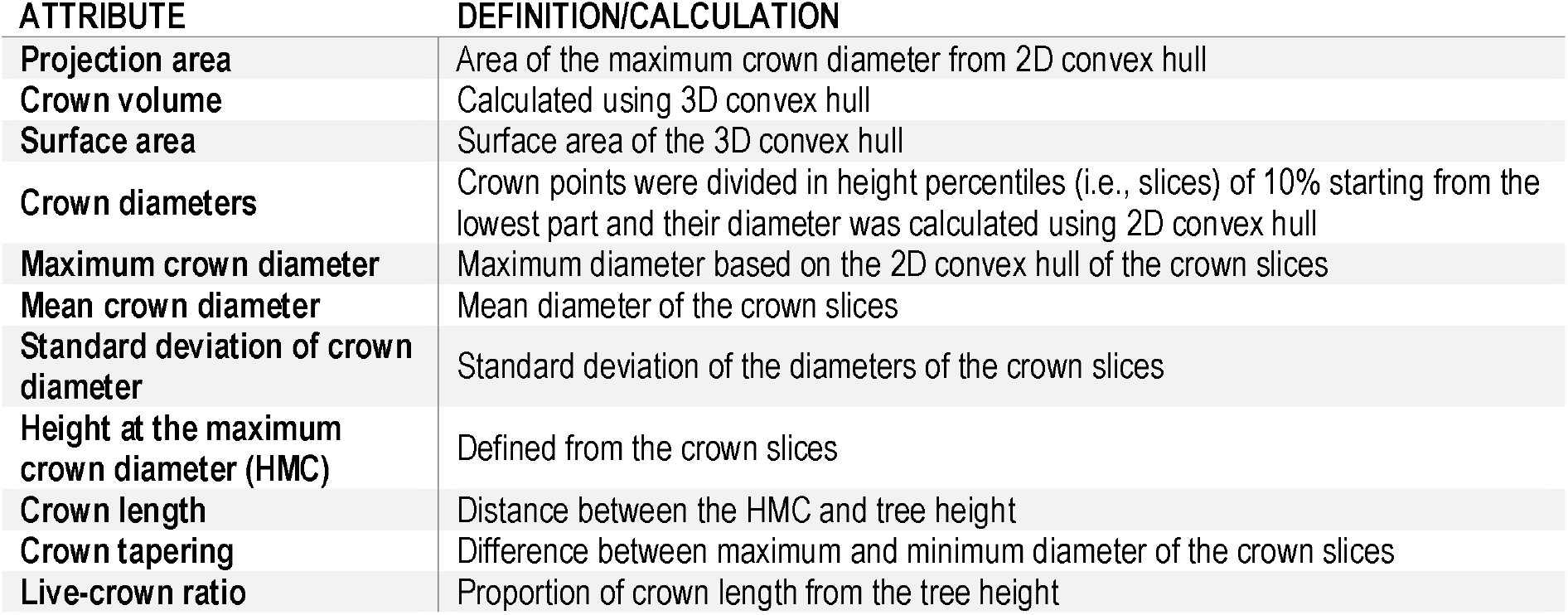
Crown attributes

Height of the maximum crown diameter (HMC) from TLS was used to define crown length (i.e., live crown base height was deducted from tree height) and live-crown ratio (i.e., proportion of crown length from tree height). Finally, stem diameter at the HMC was obtained from the taper curve and stem area at the height of the maximum crown diameter (SAHMC) was calculated as pi/4*d^2^.

Stem attributes included DBH, stem volume, height-DBH ratio (i.e., height/DBH), and cumulative volume. Tree height was obtained using the height of the highest TLS point of each tree (i.e., normalized above ground) whereas DBH was defined from taper curve obtained with a combination of circle fitting to original stem points and fitting a cubic spline (see Yrttimaa et al. 2019, Saarinen et al. 2020). Stem volume, on the other hand, was defined by considering the stem as a sequence 10 cm vertical cylinders and summing up the volumes of the cylinders using the estimated taper curve. Finally, cumulative stem volume was calculated as the height at which 50% of stem volume was accumulated.

As TLS data were only available for one time point, insitu measurements were utilized for obtaining growth information of individual Scots pine trees. Growth of DBH, tree height, stem volume, and change in height/DBH were calculated using in-situ measurements conducted in 2005-2006 (i.e., at the time of establishment of the study sites) and 2018-2019 (i.e., the latest in-situ measurements) for all live Scots pine trees that were identified from the sample plots during the latest field measurements.

### Effects of thinning on stem area at the height of the maximum crown diameter

Due to the data structure (i.e., several sample plots in each study site), a nested two-level linear mixed-effects model (Equation 1) was fitted using Restricted Maximum Likelihood included in package nlme (Pinheiro et al. 2020) of the R-software to assess the effects of thinning treatment on SAHMC.

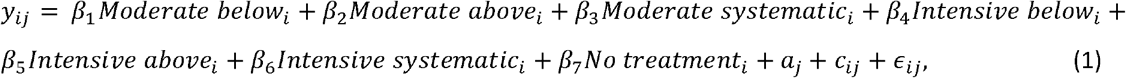

where *y* _*ij*_ is SAHMC, *β*_1_ , … *β*_2_ are fixed parameters, i, i = 1, …, M, refers to study site, j, j = 1, …, *n*_*i*_, to a plot, *a* _*j*_ and *c* _*ij*_ are normally distributed random effects for sample plot *j* and for sample plot *j* within study site *i*, respectively, with mean zero and unknown, unrestricted variance-covariance matrix, and *ϵ* _*ij*_ is a residual error with a mean zero and unknown variance. The random effects are independent across study sites and sample plots as well as residual errors are independent across trees. The effects of a study site and a sample plot within the study sites SAHMC were assessed through their variances.

### Relationship between basal area at the height of the maximum crown diameter and crown, stem, and growth attributes

Correlations between dependent and independent variables was investigated using Spearman rho rank-based correlation coefficient. Furthermore, the significance level of the correlation was investigated. The nested-two-level linear mixed-effect model in Equation 1 was utilized in investigating the possible relationship between SAHMC and different crown, stem, and growth attributes. Each crown (Table 2), stem (i.e., DBH, stem volume, height/DBH), and growth (ΔDBH, Δtree height, Δstem volume, Δheight/DBH) attribute was independently added to the Equation 1 as a predictor variable.

## Results

### The effects of stem density on crown architecture

Difference in stem density/ha varied from 430 to 470 between moderate and intensive thinning and from 310 to 960 stem/ha between no treatment and thinned (i.e. all other) plots. When thinning intensity increased (i.e. stem density/ha decreased) from moderate to intensive thinning from below, crown volume, projection area, and maximum and mean diameter increased (Figure 2) statistically significantly (p<0.05). Similarly, live-crown ratio as well as crown diameter at the bottom of a crown (i.e. 10-30 percentiles) (Figure 3) statistically significantly (p<0.05) increased when thinning intensity increased, but this was true for all thinning types. However, there was no statistically significant (p>0.05) difference in crown attributes between moderate thinnings and no treatment.

**Figure 2.**
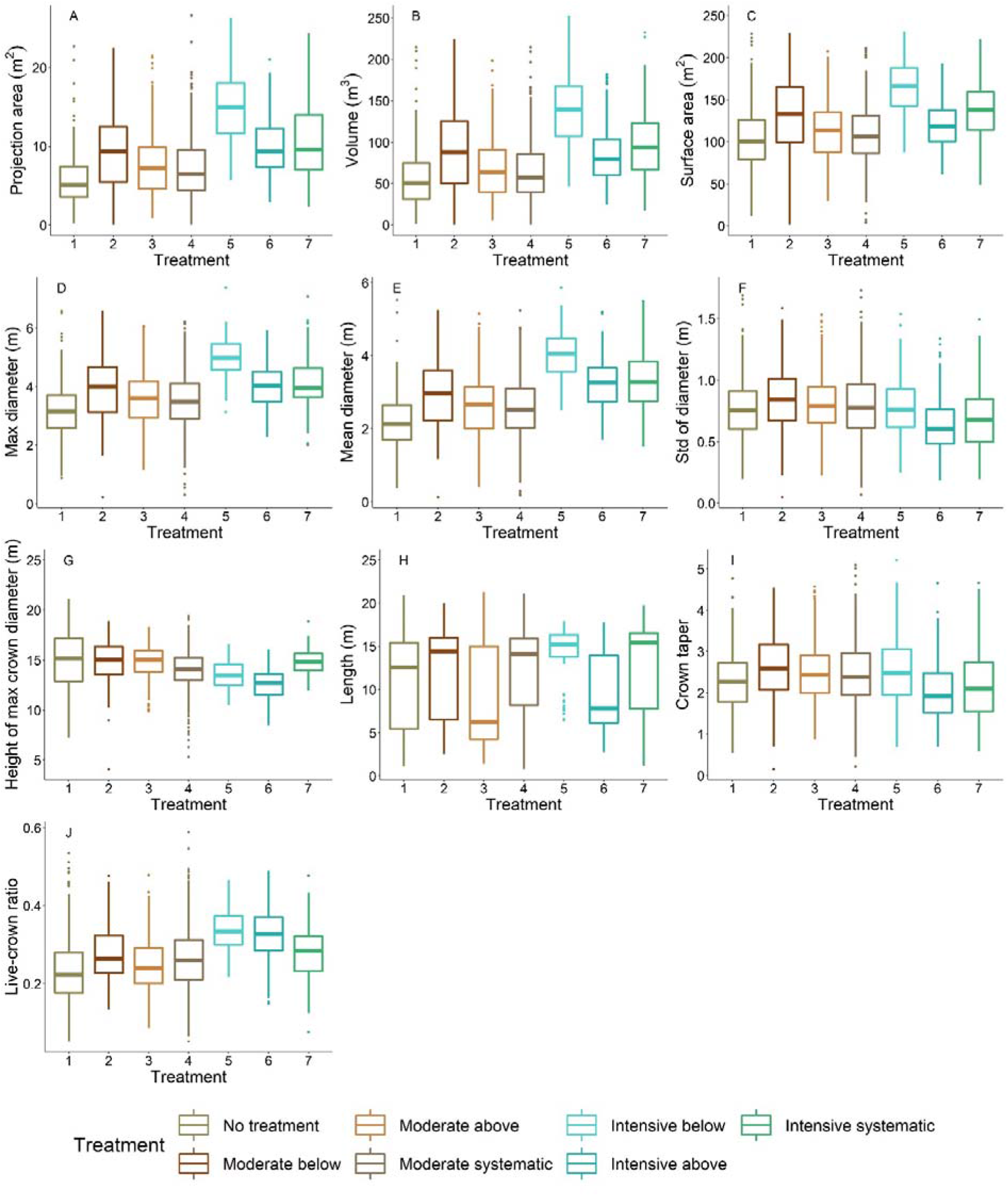
Variation of crown attributes between thinning treatments. 1 = No treatment (i.e., control), 2 = Moderate thinning from below, 3 = Moderate thinning from above, 4 = Moderate systematic thinning from above, 5 = Intensive thinning from below, 6 = Intensive thinning from above, and 7 = Intensive systematic thinning from above.

**Figure 3.**
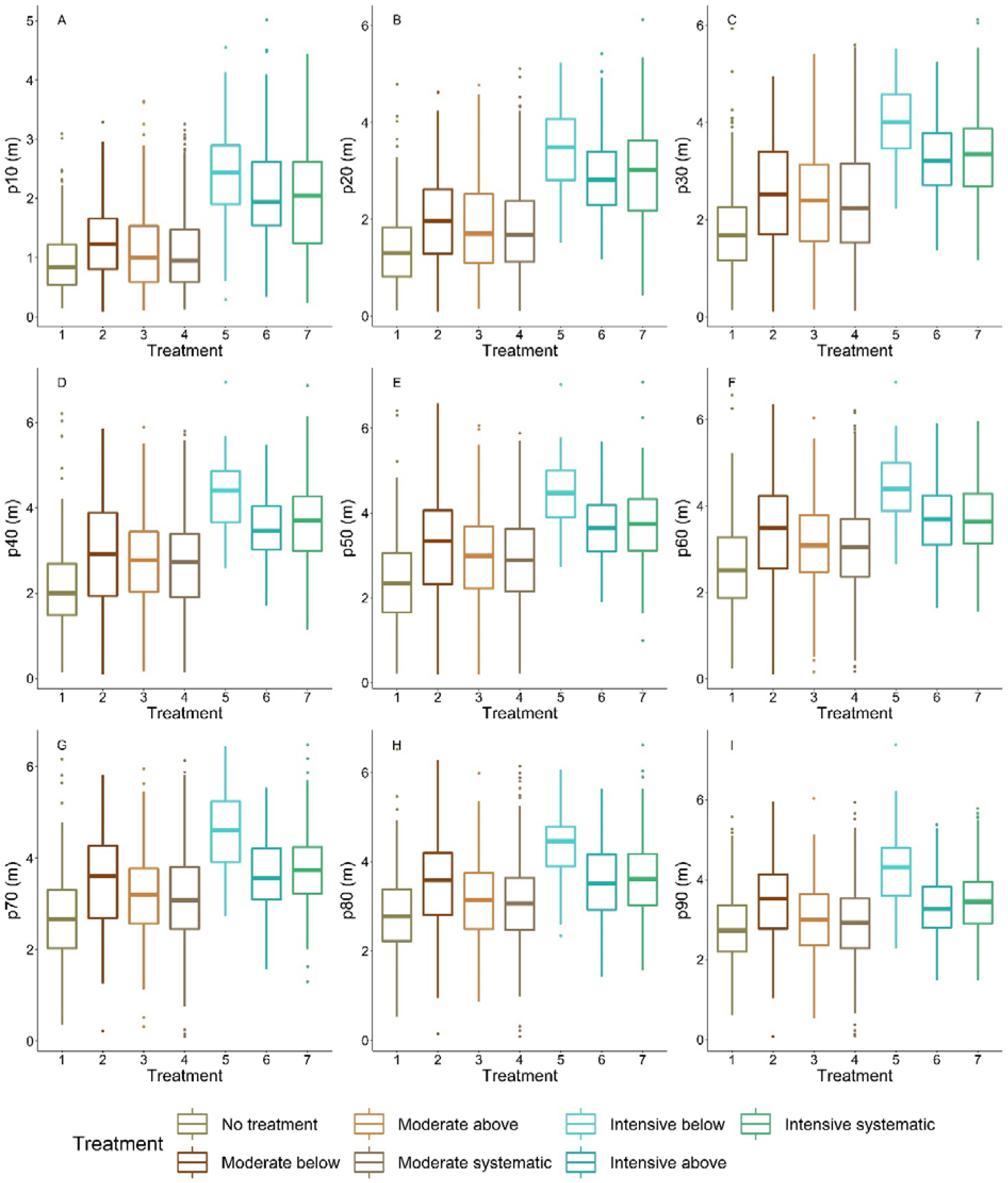
Variation of crown diameter at height percentiles between thinning treatments. P10 indicates the lowest height percentile (i.e. the most bottom part of a crown), whereas p100 is the highest height percentile (i.e. the highest part of a crown). 1 = No treatment (i.e., control), 2 = Moderate thinning from below, 3 = Moderate thinning from above, 4 = Moderate systematic thinning from above, 5 = Intensive thinning from below, 6 = Intensive thinning from above, and 7 = Intensive systematic thinning from above.

Thinning type (i.e., removal of suppressed and co-dominant or dominant trees) had a less clear effect on crown size and shape. Statistically significant (p<0.05) differences were only present in crown volume, surface and projection area, maximum and mean diameter, as well as diameters at the top part of a crown when intensive thinning from below was compared with other intensive thinnings (difference in stem density/ha between 20 and 180). In other words, in intensive thinnings crown attributes were larger when suppressed and co-dominant trees had been removed (i.e. thinning from below) compared to when dominant trees were removed (i.e. thinning from above and systematic thinning). This is also visible for example trees from different thinning treatments (Figure 4).

**Figure 4.**
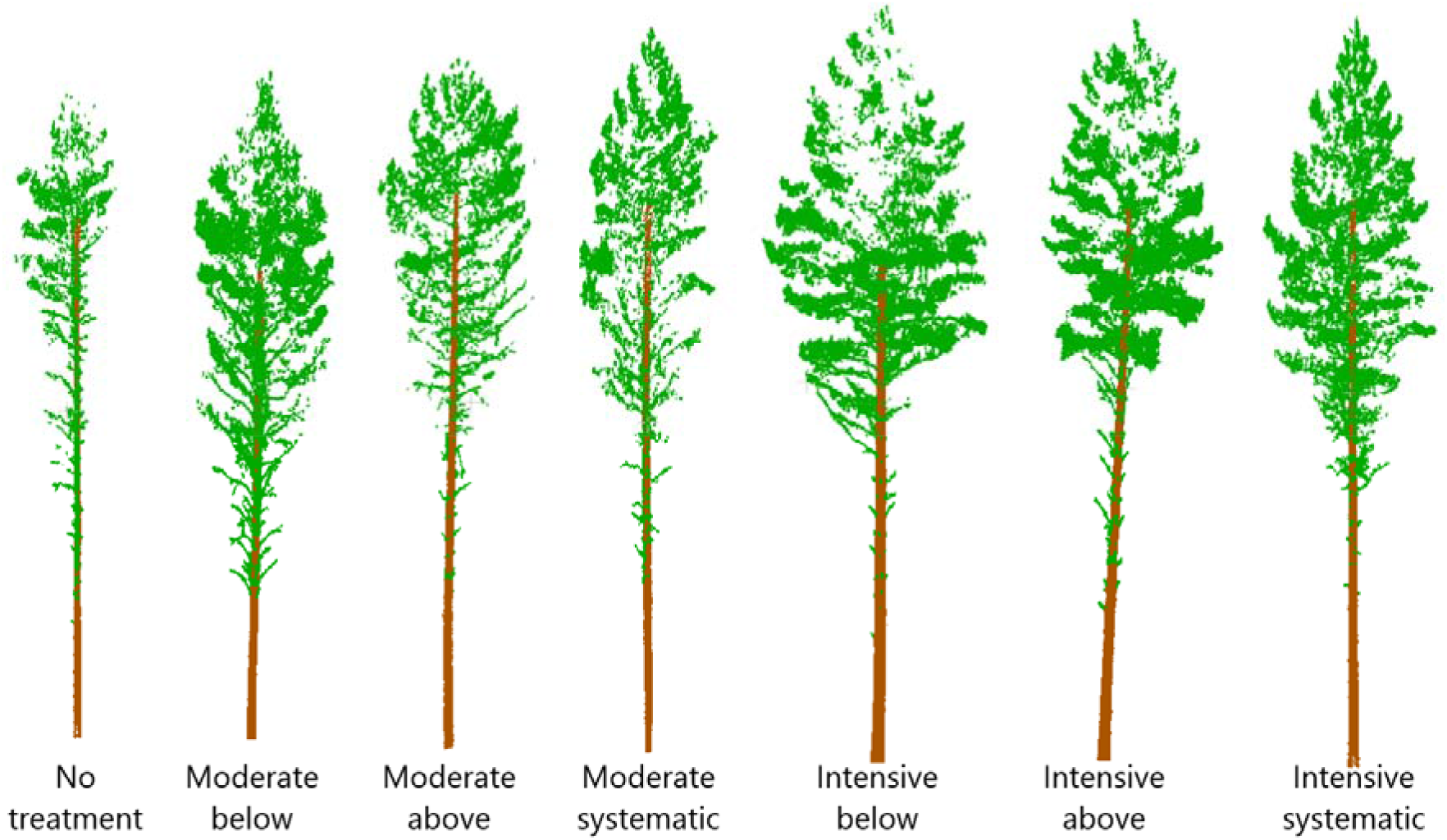
Point clouds from example trees from different thinning treatments. Stem densities of the treatments were on average ~1250, 720, 910, 940, 290, 450, and 470 stems/ha for no treatment, moderate below, moderate above, moderate systematic, intensive below, intensive above, and intensive systematic, respectively.

### The effects of stem density on stem area at the height of maximum crown diameter

SAHMC ranged from 67.4 cm^2^ to 170.2 cm^2^ being the smallest with no treatment and the largest with intensive thinning from below (Figure 5). For moderate thinnings, SAHMC was 90.6 cm^2^, on average, whereas with intensive thinnings it was 132.2 cm^2^. Lower stem densities increased SAHMC, and SAHMC was statistically significantly (p<0.05) greater when stem density increased from ~290 stems/ha (i.e. intensive below) to at least ~720 stems/ha (i.e. moderate below). In other words, SAHMC was statistically significantly different between intensive thinning from below and all other thinning treatments, including no treatment, except between intensive thinning from above.

**Figure 5.**
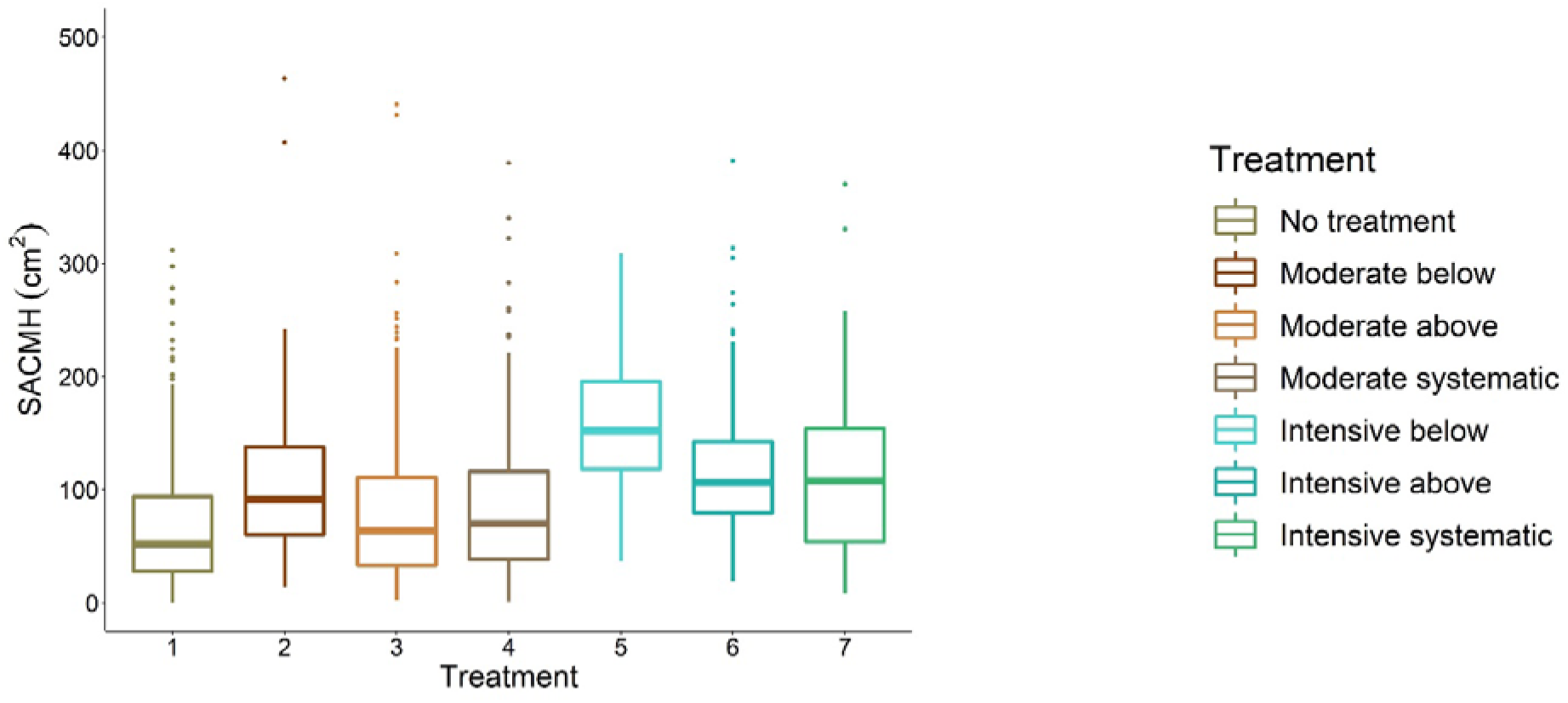
Stem area at the height of the maximum crown diameter (SAHMC) between thinning treatments. 1 = No treatment (i.e., control), 2 = Moderate thinning from below, 3 = Moderate thinning from above, 4 = Moderate systematic thinning from above, 5 = Intensive thinning from below, 6 = Intensive thinning from above, and 7 = Intensive systematic thinning from above.

### Relationship between stem area at the height of maximum crown diameter and crow and stem attributes as well as tree growth

There was high correlation (≥|0.5|) between SAHMC and most of the crown, stem, and growth attributes (Table 5). Especially, attributes characterizing crown size (i.e., projection area, volume, surface area, maximum and mean crown diameter, and live-crown ratio) and stem size (i.e. DBH, stem volume, and height at which 50% of stem volume accumulated), and size growth (i.e. DBH growth and stem volume growth) showed high positive correlation. Height/DBH ratio, on the other hand, showed negative correlation with SAHMC. Correlations between SAHMC and all crown, stem, and growth attributes were statistically significant.

**Table 5.**
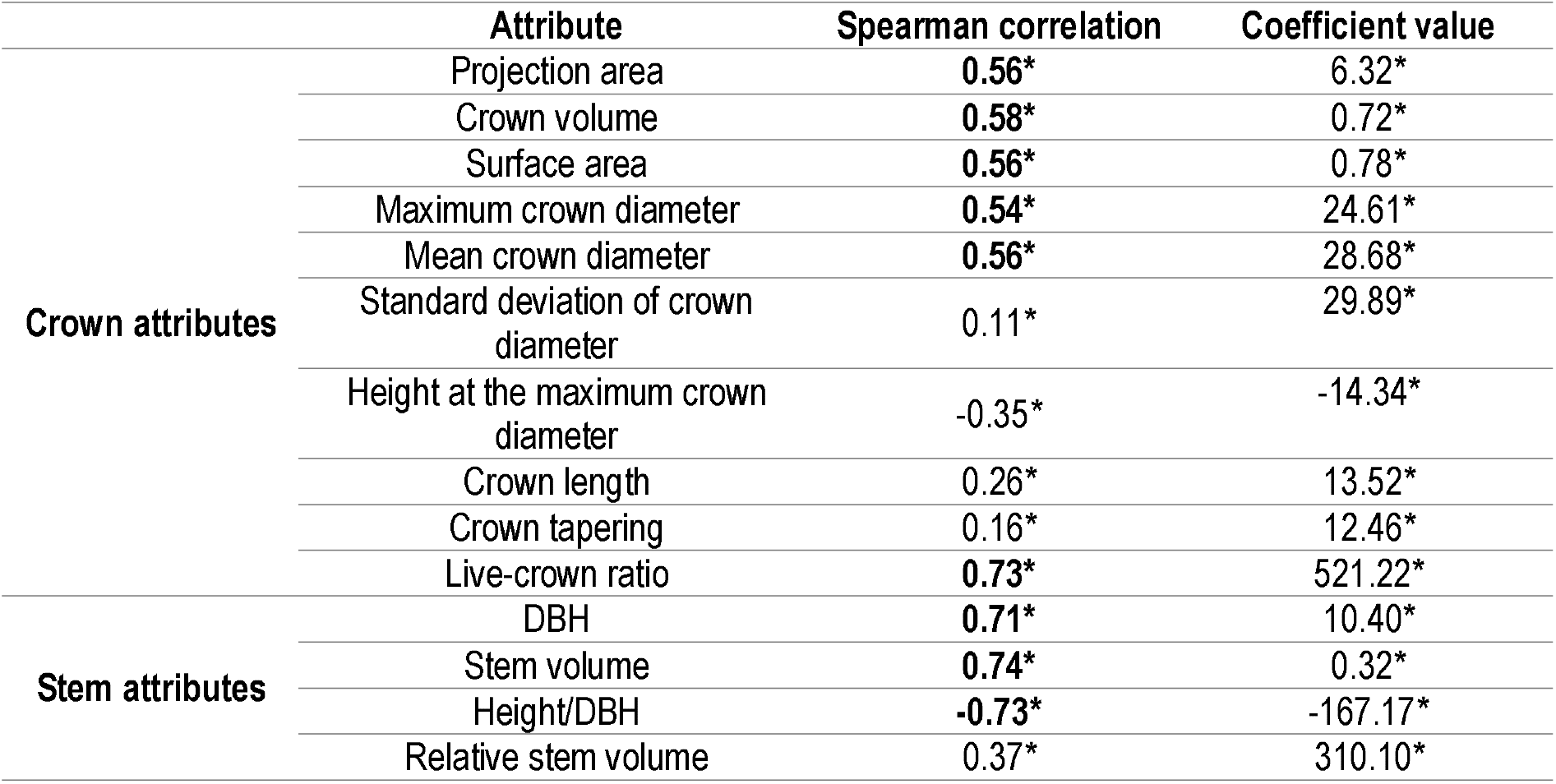
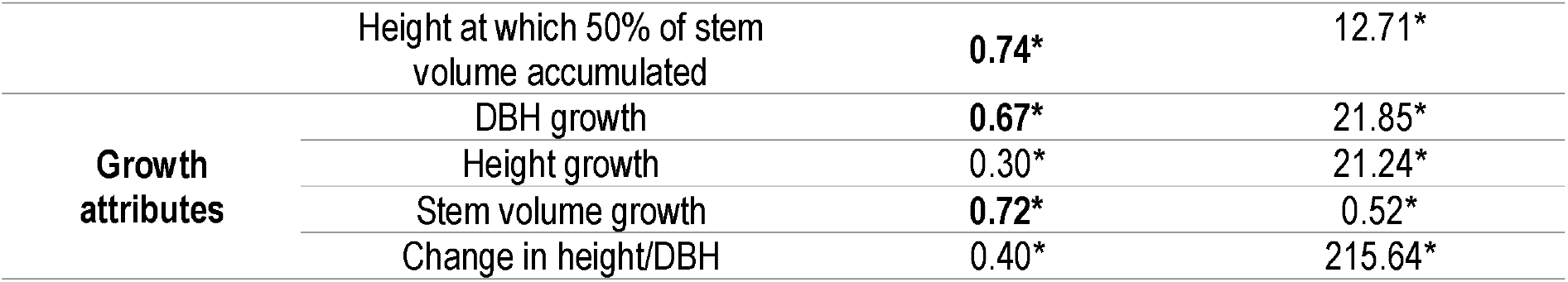
Spearman correlations between stem area at the height of the maximum crown diameter and crown, stem, and growth attributes as well as coefficient value from the nested-two-level linear mixed-effect models when each attribute was independently included as a predictor variable against stem area at the height of the maximum crown diameter. DBH = diameter at breast height. * denotes statistically significant correlation or importance in the model.

|  | Attribute | Spearman correlation | Coefficient value |
| --- | --- | --- | --- |
| **Crown attributes** | Projection area | 0.56* | 6.32* |
|  | Crown volume | 0.58* | 0.72* |
|  | Surface area | 0.56* | 0.78* |
|  | Maximum crown diameter | 0.54* | 24.61* |
|  | Mean crown diameter | 0.56* | 28.68* |
|  | Standard deviation of crown diameter | 0.11* | 29.89* |
|  | Height at the maximum crown diameter | -0.35* | -14.34* |
|  | Crown length | 0.26* | 13.52* |
|  | Crown tapering | 0.16* | 12.46* |
|  | Live-crown ratio | 0.73* | 521.22* |
| **Stem attributes** | DBH | 0.71* | 10.40* |
|  | Stem volume | 0.74* | 0.32* |
|  | Height/DBH | -0.73* | -167.17* |
|  | Relative stem volume | 0.37* | 310.10* |
|  | Height at which 50% of stem volume accumulated | **0.74\*** | 12.71* |
| **Growth attributes** | DBH growth | **0.67\*** | 21.85* |
|  | Height growth | 0.30* | 21.24* |
|  | Stem volume growth | **0.72\*** | 0.52* |
|  | Change in height/DBH | 0.40* | 215.64* |

Crown diameters at different heights also showed positive correlation (≥0.5) with SAHMC. Furthermore, the results from the nested-two-level linear mixed-effect model showed that increment in most of the crown, stem, and growth attribute, when independently included as a predictor variable, increased SAHMC. HMC and height/DBH were exceptions as their increment decreased SAHMC. Increasing live-crown ratio, relative stem volume, and change in height/DBH increased SAHMC ten times more than other crown, stem, and growth attributes, whereas the effect of increasing height/DBH was of similar magnitude but to different direction, in other words it decreased SAHMC. When each of the crown, stem, and growth attribute was separately added as a predicter variable to estimate SAHMC all of them were statistically significant (p<0.001) for the model (Table 5).

## Discussion

The results showed how thinning treatments carried out >10 years ago affected crown shape and size of Scots pine trees. As stem density decreased, crown volume, surface area, and maximum diameter increased. Also, diameter of the lower part of a crown (<80^th^ height percentile) increased with decreasing stem density. These results suggest that stem density affects crown shape and size of Scots pine trees in boreal forests. Lower stem densities (i.e. ≤700 stems/ha) also increased SAHMC. Furthermore, when crown and stem size as well as stem growth increased, also SAHMC grew.

One of the traditional parameters used for characterizing crown architecture is live-crown ratio and the results here showed that it differed between stem densities, similarly to the findings by Kellomäki & Tuimala (1981). Additionally, there were significant differences between more advanced parameters (namely crown surface area and volume) at least amongst the sparsest stem densities (i.e., intensive thinning). Finally, the study confirmed the results presented by Oker-Blom & Kellomäki (1982) as the lowest part of a Scots pine tree crown was larger in low stem densities. Thus, the use of TLS for obtaining enhanced information on canopy structure and architecture can be justified.

There is uncertainty in the SAHMC as the HMC may not represent the height of crown base height, which is traditionally used for crown length and live-crown ratio. Thus, also SAHMC may not represent the true dcb. However, it has not been traditionally feasible to measure dcb from standing trees, whereas measurements on stem diameters from TLS data offer this. Thus, our results show a way towards assessing the usefulness of dcb as a proxy for growth potential of individual trees. There was strong correlation (≥0.5) between SAHMC and most of the crown attributes (e.g. crown volume, surface area, diameter, and live-crown ratio) but also with DBH and stem volume, and their growth. This indicates, that dcb or SAHMC could also be used when assessing growth potential, and TLS offers a means for obtaining this information.

Studies utilizing TLS in assessing tree development include European beech (*Fagus sylvatica* [L.]) (Juchheim et al. 2017, Georgi et al. 2018) and holm oak (*Quercus ilex* L.) (Bogdanovich et al. 2021). Juchheim et al. (2017) found that increasing thinning intensity increased crown surface area of European beech, which is in line with our results for Scots pine. Georgi et al. (2018) reported that crown size (i.e. crown volume, projection area, surface area, length, and live-crown ratio) of European beech trees growing in stands without forest management in ≥50 years was statistically significantly lower compared to European beech trees growing in managed stands or stands with ≤20 years without forest management. Our results showed that only intensive thinning resulted in statistically significant difference in crown attributes (e.g. crown volume, projection area, and maximum and mean diameter) when compared with moderate thinning and no treatment. In other words, moderate thinning had no effect on crown size when compared with no treatment.

As height/DBH and absolute height of the crown base have been identified as indicators for tree vitality (Longuetaud et al. 2006), this study presented a means for obtaining those attributes. Height/DBH has been shown to increase as forest management intensity increased (Saarinen et al. 2020), whereas HMC did not differ significantly between tree densities in this study. However, this study provided dcb and stem cross-sectional area at the HMC which enables studies on their suitability as proxies for growth potential.

This study concentrated on investigating crown structure of individual Scots pine trees in different stem densities. Increasing stem density decreased crown size, confirming our hypothesis (H1). With low stem densities (i.e., intensive thinning), crown size also increased when suppressed and co-dominant trees were removed (i.e., thinning from below) partly confirming the H2 (i.e., no difference in moderate thinnings). Furthermore, a relationship between SAHMC and crown and stem attributes was found. Thus, this study showed how tree density affects crown shape and size of Scots pine trees and how they are adapted to the growing conditions of the trees. As stem density can be regulated through forest operations such as thinning, the results of this study can be utilized when planning management actions.

## Conclusions

Stem densities affected crown size and shape of Scots pine trees growing in boreal forests. When growing in a denser forest, the crown size of Scots pine tree decreased, indicating more competition on light between adjacent tree crowns. Although this has been known for decades as growth and yield studies have a long history, this study provided quantitative attributes assessing crown size (e.g. crown volume, projection area, surface area, diameter) and shape (i.e. diameters at different heights of a crown, their mean and standard deviation) of Scots pine trees. Additionally, the study provided stem diameter and cross-sectional area at the height of maximum crown diameter (i.e. SAHMC) that can be assumed to present crown base height. Increasing forest management intensity increased the SAHMC and there was strong relationship between it and crown, stem, and growth attributes. Thus, it can be concluded that this study expanded our knowledge on the crown architecture of Scots pine trees of different size growing in different conditions (i.e., different stem densities) that were a result of past forest management activities. This was enabled with detailed 3D TLS data that offered quantitative and more comprehensive characterization of Scots pine crowns and growth potential.

## Acknowledgements

The study was funded by the Academy of Finland postdoctoral projects 315079, 345166, 330422 as well as Finnish Flagship Programme of the Academy of Finland (grant numbers 337127, 337655, 337656).

## References

Aaltonen, V.T. 1925. Metsikön itseharvenemisesta ja puiden kasvutilasta luonnonmetsissä. Communicationes Ex Instituto Quaestionum Forestalium Finlandiae 9: 1–17. [Ber der Selbstabscdeidung un den Wuchsraum de Bäume in Naturbestanden]. In Finnish with German summary.

Barbeito I., Dassot, M., Bayer, D., Collet, C., Drössler, L., Löf, M., del Rio, M., Ruiz-Peinado, R., Forrester, D.I., Bravo-Oviedo, A., Pretzsch, H. 2017. Terrestrial laser scanning reveals differences in crown structure of *Fagus sylvatica* in mixed *vs.* pure European forests. Forest Ecology and Management 405: 381–390. http://dx.doi.org/10.1016/j.foreco.2017.09.043

Bauhus, J., van Winden, A.P., Nicotra, A.B. 2004. Aboveground interactions and productivity in mixed.species plantations of *Acacia mearnsii* and *Eucalyptus globulus*. Canadian Journal of Forest Research 34: 686–694. https://doi.org/10.1139/x03-243

Bayer, D., Seifert, S., Pretzsch, H. 2013. Structural crown properties of Norway spruce (*Picea abies* [L.] Karst.) and European beech (*Fagus sylvatica* [L.]) in mixed versus pure stands revealed by terrestrial laser scanning. Trees 27: 1035–1047. https://doi.org/10.1007/s00468-013-0854-4

Binkley, D., Camargo Campoe, O., Gspaltl, M., Forrester, D.I. 2013. Light absorption and use efficiency in forests: Why patterns differ for trees and stands. Forest Ecology and Management 288: 5–13. https://doi.org/10.1016/j.foreco.2011.11.002

Bogdanovich E., Perez-Priego, O., El-Madany, T.S., Guderle, M., Pacheco-Labrador, J., Levick, S.R., Moreno, G., Carrara, A., Martín, M.P., Migliavacca, M. 2021. Using terrestrial laser scanning for characterizing tree structural parameters and their changes under different management in a Mediterranean open woodland. Forest Ecology and Management 486: 118945. https://doi.org/10.1016/j.foreco.2021.118945

Chianucci, F., Puletti, N., Grotti, M., Ferrara, C., Giorcelli, A., Coaloa, D., Tattoni, C. 2020. Nondestructive tree stem and crown volume allometry in hybrid poplar plantations derived from terrestrial laser scanning. Forest Science 66: 737–746. https://doi.org/10.1093/forsci/fxaa021

Curtin, R.A. 1970. Dynamics of tree and crown structure in *Eucalytus obliqua*. Forest Science 16: 321–328. https://doi.org/10.1093/forestscience/16.3.321

Dieler, J. & Pretzsch, H. 2013. Morphological plasticity of European beech (*Fagus sylvatica* L.) in pure and mixed-species stands. Forest Ecology and Management 295: 97–108. https://doi.org/10.1016/j.foreco.2012.12.049

Fernández-Sarría, A., Velázquez-Marí, B., Sadjak, M-. Mart’nez, L., Estornell, J. 2013. Residual biomass calculation from individual tree architecture using terrestrial laser scanner and ground-level measurements. Computers and Electronics in Agriculture 93: 90–97. https://doi.org/10.1016/j.compag.2013.01.012

Flower-Ellis, J., Albrektsson, A., Olsson, L. 1976. Structure and growth of some young Scots pine stands: (1) dimensional and numerical relationships. Swedish Conifer Project. Technical Report 3: 1–98.

Forrester, D.I. 2014. The spatial and temporal dynamics of species interactions in mixed-species forests: From pattern to process. Forest Ecology and Management 312: 282–292. https://doi.org/10.1016/j.foreco.2011.11.002

Georgi, L., Kunz, M., Fichter, A., Härdtle, W., Reich, K.R., Strum, K., Welle, T., von Oheimb, G. 2018. Long-term abandonment of forest management has a strong impact on tree morphology and wood volume allocation pattern of European beech (*Fagus sylvatica* L.). Forests 9: 704. https://doi.org/10.3390/f9110704

Georgi, L., Kz, M., Fichtner, A., Reich, K.F., Bienert, A., Maas, H.-G., von Oheimb, G. 2021 Effects of local neighbourhood diversity on crown structure and productivity of individual tree in mature mixed-species forests. Forest Ecosystems 8: 26. https://doi.org/10.1186/s40663-021-00306-y

Grinrich, S.F. 1967. Measuring and evaluating stocking and stand density in upland hardwood forests in the central states. Forest Science 13:38–53. https://doi.org/10.1093/forestscience/13.1.38

Hashimoto, R. 1986. Comparative study of thinning methods in young sugi (*Cryptomeria japonica*) plantations: changes in canopy structure and light environment. In: Fujimori, T., Whitehead, D. (Eds) Crown and canopy structure in relation to productivity. Forestry and Forest Products Research Institute, Ibaraki, Japan

Heikinheimo, O. 1953. Puun rungon luontaisesta karsiutumisesta. Communtiones Instituti Forestalis Fenniae 41(5): 1–39. [On natural pruning of tree stems]. In Finnish with English summary.

Hess, C., Härdtle, W., Kunz, M., Fichtner, A., von Oheimb, G. 2018. A high-resolution approach for the spatiotemporal analysis of forest canopy space using terrestrial laser scanning. Ecology and Evolution 8: 6800–6811. https://doi.org/10.1002/ece3.4193

Hildebrand, M., Perles-Garcia, M.D., Kunz, M., Härdtle, W., von Oheimb, G., Fichter, A. 2021. Tree-tree interactions and crown complementarity: The role of functional diversity and branch traits for canopy acking. Basic and Applied Ecology 50: 217–227. https://doi.org/10.1016/j.baae.2020.12.003

Hu, M., Lehtonen, A., Minunno, F., Mäkelä, A. 2020. Age effect on tree structure and biomass allocation in Scots pine (*Pinus sylvestris* L.) and Norway spruce (*Picea abies* [L.] Karst.). Annals of Forest Science 77: 90. https://doi.org/10.1007/s13595-020-00988-4

Hynynen, J. 1995. Predicting tree crown ratio for unthinned and thinned Scots pine stands. Canadian Journal of Forest Research 25:57–62. https://doi.org/10.1139/x95-007

Ilomaki, S., Nikinmaa, E. &Makela, A. (2003). Crown rise due to competition drives biomass allocation in silver birch. Canadian Journal of Forest Research 33: 2395–2404. https://doi.org/10.1139/x03-164

Juchheim, J., Annighöfer, P., Ammer, C., Calders, K., Raumonen, P., Seidel, D. 2017. How management intensity and neighborhood composition affect the structure of beech (*Fagus sylvatics* L.) trees. Trees 31: 1723–1735. https://doi.org/10.1007/s00468-017-1581-z

Juchheim, J., Ehbrecht, M., Schall, P., Ammer, C., Seidel, D. 2019. Effect of species mixing on stand structural complexity. Forestry 93: 75–83. https://doi.org/10.1093/forestry/cpz046

Kantola, A. & Mäkelä, A. (2004). Crown development in Norway spruce [Picea abies (L.) Karst.]. Trees 18: 408–421. https://doi.org/10.1007/s00468-004-0319-x

Kellomäki, S. 1980. Growth dynamics of young Scots pine crowns. Communicationes Instituti Forestalis Fenniae 98(4): 1–50.

Kellomäki, S., Tuimala, A. 1981. Puuston tiheyden vaikutus puiden oksikkuuteen taimikko- ja riukuvaiheen männiköissä. Folia Forestalia 478: 1–27. [Effect of stand density on branchiness of young Scot pines]. In Finnish with English summary.

Krajicek, J.E., Brinkman, K.A., Gingrich, S.F. 1961. Crown Competition—A Measure of Density. Forest Science 7: 35–42. https://doi.org/10.1093/forestscience/7.1.35

Larson, P.R. 1963. Stem form development of forest trees. Forest Science 9: 1–42. https://doi.org/10.1093/forestscience/9.s2.a0001

Lehnebach, R., Beyer, R., Letort, V., Heuret, P. 2018. The pipe model theory half a century on: a review. Annals of Botany 121: 773–795. https://doi.org/10.1093/aob/mcx194

Lehtonen, A., Heikkinen, J., Petersson, H., Ťupek, B., Liski, E., Mäkelä, A. 2019. Scots pine and Norway spruce foliage biomass in Finland and Sweden – testing traditional models vs. the pipe model theory. Canadian Journal of Forest Research 50: 146–154. https://doi.org/10.1139/cjfr-2019-0211

Liang, X., Kankare, V., Yu, X., Hyyppä, J., Holopainen, M. 2014. Automated stem curve measurement using terrestrial laser scanning. IEEE Transactions on Geoscience and Remote Sensing 52: 1739–1748. https://doi.org/10.1109/TGRS.2013.2253783

Longuetaud, F., Mothe, F., Leban, J.-M., Mäkelä, A. 2006 *Picea abies* sapwood width: Variations within and between trees. Scandinavian Journal of Forest Research 21: 41–53. https://doi.org/10.1080/02827580500518632

Mäkinen, H., Isomäki, A. 2004. Thinning intensity and long-term changes in increment and stem form of Scots pine trees. Forest Ecology and Management 201(1-3): 21–34. https://doi.org/10.1016/j.foreco.2004.07.028

Martin-Ducup, O., Schneider, R., Fournier, R.A. 2016. Response of sugar maple (*Acer saccharum*, Marsh.) tree crown structure to competition in pure versus mixed stands. Forest Ecology and Management 374: 20–32. http://dx.doi.org/10.1016/j.foreco.2016.04.047

Metz, J., Seidel, D., Schall, P., Scheffer, D., Schulze, E.-D-. Ammer, C. 2013. Crown modeling by terrestrial laser scanning as an approach to assess the effect of aboveground intra- and interspecific competition on tree growth. Forest Ecology and Management 213: 275–288. http://dx.doi.org/10.1016/j.foreco.2013.08.014

Nikinmaa E. 1992. Analyses of the growth of Scots pine: matching structure with function. Acta Forestalia Fennica 235: 7681. https://doi.org/10.14214/aff.7681

Nykänen, M.L., Peltola, H., Quine, C.P., Kellomäki, S., Broadgate, M. 1997. Factors affecting snow damage of trees with particular reference to European conditions. Silva Fennica 31(2): 193–213. https://doi.org/10.14214/sf.a8519

Oker-Blom, P., Kellomäki, S. 1982. Metsikön tiheyden vaikutus puun latvuksen sisäiseen valoilmastoon ja oksien kuolemiseen – Teoreettinen tutkimus. Folia Forestalia 509: 1–14. [Effect of stand density on the within-crown light regime and dying-off of branches – Theoretical study]. In Finnish.

Owen, H.J.F., Flynn, W.R.M., Lines, E.R. 2021. Competitive drivers of interspecific deviations of crown morphology from theoretical predictions measured with terrestrial laser scanning. Journal of Ecology 109: 2612–2628. https://doi.org/10.1111/1365-2745.13670

Pretzsch, H. 2014. Canopy space filling and tree crown morphology in mixed-species stands compared with monocultures. Forest Ecology and Management 327: 251–264. https://doi.org/10.1016/j.foreco.2014.04.027

Pretzsch, H. 2019. The effect of tree crown allometry on community dynamics on mixed-species stands versus monocultures. A review and perspectives for modeling and silvicultural regulation. Forests 10: 810. https://doi.org/10.3390/f10090810

Pretzsch, H., Biber, P., Uhl, E., Dahlhausen, J., Rötzer, T., Calderntey, J., Koike, T., van Con, T., Chavanne, A., Seifert, T., du Toit, B., Farnden, C., Pauleit, S. 2015. Crown size and growing space requirement of common tree species in urban centres, parks, and forests. Urban Forestry & Urban Greening 14: 466–479. https://doi.org/10.1016/j.ufug.2015.04.006

Pyörälä, J., Liang, X., Saarinen, N., Kankare, V., Wang, Y., Holopainen, M., Hyyppä, J., Vastaranta, M. 2018a. Assessing branching structure for biomass and wood quality estimation using terrestrial laser scanning point clouds. Canadian Journal of Remote Sensing 44: 462–475. https://doi.org/10.1080/07038992.2018.1557040

Pyörälä, J., Liang, X., Vastaranta, M., Saarinen, N., Kankare, V., Wang, Y., Holopainen, M., Hyyppä, J. 2018b. Quantitative assessment of Scots pine (*Pinus sylvestris* L.) Whorl structure in a forest environment using terrestrial laser scanning. IEEE Journal of Selected Topics in Applied Earth Observations and Remote Sensing 11: 3598–3607. https://doi.org/10.1109/JSTARS.2018.2819598

Pyörälä, J., Saarinen, N., Kankare, V., Coops, N.C., Liang, X., Wang, Y., Holopinen, M., Hyyppä, J., Vastaranta, M. 2019. Variability of wood properties using airborne and terrestrial laser scanning. Remote Sensing of Environment 235: 111474. https://doi.org/10.1016/j.rse.2019.111474

Rais, A., Jacobs, M., van de Kuilen, J.-W.G., Pretzsch, H. 2021. Crown structure of European beech (*Fagus sylvatica*): a noncausal proxy for mechanical-physical wood properties. Canadian Journal of Forest Research 51: 834–841. https://doi.org/10.1139/cjfr-2020-0382

Saarinen, N., Kankare, V., Vastaranta, M., Luoma, V., Pyörälä, J., Tanhuanpää, T., Liang, X., Kaartinen, H., Kukko, A., Jaakkola, A., Yu, X., Holopainen, M., Hyyppä, J. 2017. Feasibility of Terrestrial Laser Scanning for Collecting Stem Volume Information from Single Trees. ISPRS Journal of Photogrammetry and Remote Sensing 123:140–158. https://doi.org/10.1016/j.isprsjprs.2016.11.012

Saarinen, N., Kankare, V., Yrttimaa, T., Viljanen, N., Honkavaara, E., Holopainen, M., Hyyppä, J., Huuskonen, S., Hynynen, J., Vastaranta, M. 2020. Assessing the effects of thinning on stem growth allocation of individual Scots pine trees. Forest Ecology and Management 474: 118344. https://doi.org/10.1016/j.foreco.2020.118344

Saarinen, N., Calders, K., Kankare, V., Yrttimaa, T., Junttila, S., Luoma, V., Huuskonen, S., Hynynen, J., Verbeeck, H. 2021. Understanding 3D structural complexity of individual Scots pine trees with different management history. Ecology and Evolution 11(6): 2561–2572. https://doi.org/10.1002/ece3.7216

Seidel, D., Leuschner, C., Müller, A., Krause, B. 2011. Crown plasticity in mixed forests-Quantifying asymmetry as a measure of competition using terrestrial laser scanning. Forest Ecology and Management 261: 2123–2132. https://doi.org/10.1016/j.foreco.2011.03.008

Seidel, D., Scahll, P., Gille, M., Ammer, C. 2015. Relationship between tree growth and physical dimensions of *Fagus sylvatica* crown assessed from terrestrial laser scanning. iForest Biosciences and Forestry 8: 735–742. https://doi.org/10.3832/ifor1566-008

Seidel, D., Annighöfer, P., Stiers, M., Zemp, C.D., Burkardt, K., Ehbrecht, M., Willim, K., Kreft, H., Hölscher, D., Ammer, C. 2019a. How a measure of tree structural complexity relates to architectural benefit◻to◻cost ratio, light availability, and growth of trees. Ecology and Evolution 9: 7134–7142. https://doi.org/10.1002/ece3.5281

Seidel, D., Ehbrecht, M. Dorji, Y., Jambay, J., Ammer, C., Annighöfer, P. 2019b. Identifying architectural characteristics that determine tree structural complexity. Trees 33: 911–949. https://doi.org/10.1007/s00468-019-01827-4

Seymour, R.S. & Smith, D.M. 1987. A new stocking guide formulation applied to eastern white pine. Forest Science 33: 469–484. https://doi.org/10.1093/forestscience/33.2.469

Shinozaki K, Yoda K, Hozumi K, Kira T. 1964. A quantitative analysis of plant form – the pipe model theory. I. Basic analyses. Japanese Journal of Ecology 14: 97–105.

Valentine, H.T. & Mäkelä, A. 2005. Bridging process-based and empirical approaches to modeling tree groeth. Tree Physiology 25: 769–779. https://doi.org/10.1093/treephys/25.7.769

Valinger, E., Lundqvist, L., Bondesson, L. 1993. Assessing the risk of snow and wind damage from tree physical characteristics. Forestry 66(3): 249–260. https://doi.org/10.1093/forestry/66.3.249

Valinger, E., Lundqvist, L., Brandel, G. 1994. Wind and snow damage in a thinning and fertilisation experiment in *Pinus sylvestris*. Scandinavian Journal of Forest Research 9:129–134. https://doi.org/10.1080/02827589409382822

Vanninen, P., Ylitalo, H., Sievanen, R. & Makela, A. (1996). Effects of age and site quality on the distribution of biomass in Scots pine (*Pinus sylvestris* L.). Trees 10: 231–238. https://doi.org/10.1007/BF02185674

Yrttimaa, T., Saarinen, N., Kankare, V., Liang, X., Hyyppä, J., Holopainen, M., Vastaranta, M. 2019. Investigating the feasibility of multi-scan terrestrial laser scanning to characterize tree communities in southern boreal forests. Remote Sensing 11(12): 1423. https://doi.org/10.3390/rs11121423

Yrttimaa, T., Saarinen, N., Kankare, V., Hynynen, J., Huuskonen, S., Holopainen, M., Hyyppä, J., Vastaranta, M. 2020. Performance of terrestrial laser scanning to characterize managed Scots pine (Pinus sylvestris L.) stands is dependent on forest structural variation. ISPRS Journal of Photogrammetry and Remote Sensing 168: 277–287. https://doi.org/10.1016/j.isprsjprs.2020.08.017

Zhu, Z., Klelnn, C., Nöle, N. 2021. Assessing tree crown volume–A review. Forestry 94: 18–35. https://doi.org/10.1093/forestry/cpaa037

